# An effective *C. elegans* CRISPR training module for high school and undergraduate summer research experiences in molecular biology

**DOI:** 10.1101/2023.09.13.557573

**Authors:** Carmen Herrera Sandoval, Christopher Borchers, Scott Takeo Aoki

**Affiliations:** Department of Biochemistry and Molecular Biology; School of Medicine; Indiana University Purdue University Indianapolis; Indianapolis, IN, 46202; Indiana BioMedical Gateway (IBMG) Program; School of Medicine; Indiana University Purdue University Indianapolis; Indianapolis, IN, 46202

**Keywords:** CRISPR, gene editing, *C. elegans*, training module

## Abstract

Engaging in research experiences as a high school or undergraduate student interested in science, technology, engineering, and mathematics (STEM) is pivotal for their academic and professional development. A structured teaching framework can help cultivate a student’s curiosity and passion for learning and research. In this study, an effective eight-week training program has been created that encompasses fundamental molecular biology principles and hands-on laboratory activities. This curriculum focuses on using clustered regularly interspaced short palindromic repeats (CRISPR) gene editing in the *Caenorhabditis elegans* model organism. Through pre- and post-program assessments, substantial enhancements in students’ molecular biology proficiency and enthusiasm for scientific exploration was observed. Overall, this diligently crafted training module that employs *C. elegans* as an educational tool to instruct inexperienced students has demonstrated its accessibility and ability to engage students in molecular biology and gene editing methodologies.

## 1. INTRODUCTION

Research experiences for students interested in science, technology, engineering, and mathematics (STEM)-related fields help foster a student’s academic and professional development. Students who participate in research during the first two years of college are more likely to remain in STEM majors (Nagda *et al*. 1998) and self-report higher confidence in their science learning abilities, especially for women and historically marginalized minorities (Auchincloss *et al*. 2014; Bangera and Brownell 2014). Through research experiences, students develop critical thinking skills, gain confidence in their ability to become successful professionals (Adebisi 2022) and are more engaged with their coursework after their summer experiences (Lopatto 2007). STEM students also benefited from targeted one-on-one mentoring (Mcsweeney *et al*. 2018). Positive research experiences increase participants desires to earn a doctoral-level degree (Lessard *et al*. 2021) and contribute to their overall success in graduate school (Vincent-ruz *et al*. 2018). Thus, effective research training modules can directly improve STEM learning for all students, regardless of their academic background or career goals.

The establishment of Clustered Regularly Interspaced Short Palindromic Repeats (CRISPR) gene editing in science and popular culture opens opportunities to engage students in molecular biology concepts. CRISPR– CRISPR-associated protein (Cas) mediated genome editing is a prokaryotic mechanism for adaptive immunity against viruses and other foreign invaders (Jiang and Doudna 2017). CRISPRs were first discovered in the sequences of DNA from *Escherichia coli* (Ishino et al. 1987) and *cas* genes later shown to encode proteins with endonuclease activity (Jinek *et al*. 2012; Jiang and Doudna 2017). Currently, CRISPR-Cas has become a widespread method used in scientific laboratories and a common topic in biology curricula (Dahlberg and Groat Carmona 2018). Recombinant Cas proteins, like the *S. pyogenes* Cas9 (e.g. (Jinek *et al*. 2012)), can be combined with chemically synthesized RNAs to form an enzyme complex capable of targeted DNA cleavage **(Fig 1A)**. Cas9-mediated genome editing can be divided into three steps (Jiang and Doudna 2017): 1) DNA site recognition, 2) DNA cleavage, and 3) DNA repair. RNA directs Cas9 to the gene target sequence through complimentary base pairing (Jiang and Doudna 2017). Once paired with the specific sequence, Cas9 will cleave the DNA site, creating a double-stranded break (DSB) **(Fig 1A)** (Jiang and Doudna 2017). The DSB is repaired by the host cellular machinery (Jiang and Doudna 2017), either by error-prone nonhomologous end joining (NHEJ) (Lieber 2010) or by homology direct repair (San filippo *et al*. 2008). Through this method, genetic regions can be removed, or coding regions inserted to create null mutations, large deletions, point mutants, addition of protein or fluorescent tags, and other modifications to study the biology and pathology of the gene of interest.

**Figure 1.**
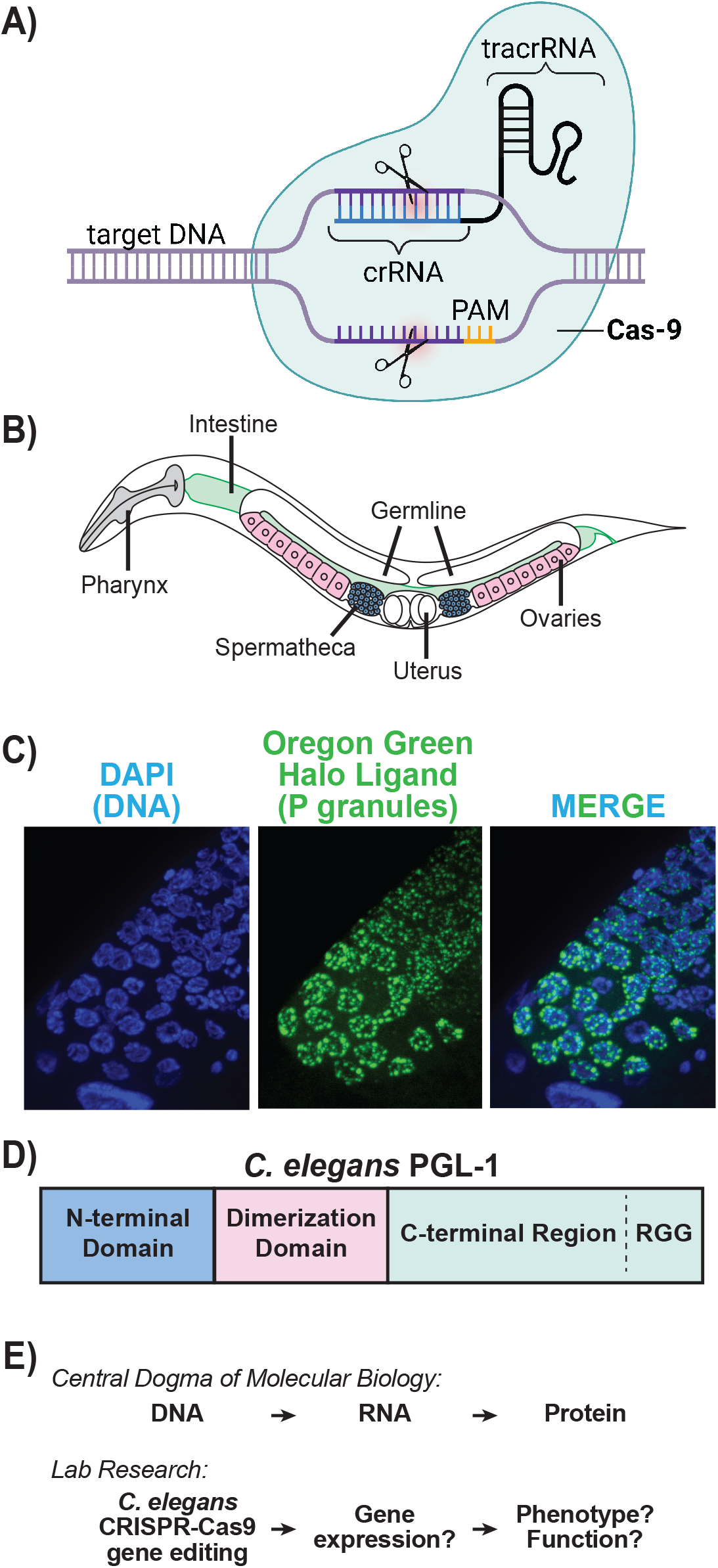
Essential concepts for summer students research experience. **(A)** Diagram of CRISPR-Cas-9 components and RNA-mediated cleavage. Trans-activating CRISPR RNA (tracrRNA) base pairs with CRISPR RNA (crRNA) to form a guide RNA. tracrRNA and crRNA interaction is crucial for target recognition and cleavage. The protospacer adjacent motif (PAM) sequence is required for Cas9 nuclease activity, causing double stranded DNA breaks 3-4 nucleotides downstream from the PAM site. The canonical PAM sequence is 5’-NGG-3’, where “N” is any nucleobase followed by two guanine (G) nucleobases. Figure created with Biorender (www.biorender.com). **(B)** *C. elegans* anatomy of the adult hermaphrodite. The CRISPR-Cas9 mix is injected into the germline directly. PGL-1 is also expressed in the germline. **(C)** Confocal microscopy of P granules in the *C. elegans* adult germline. These germ granules are found at the nuclear periphery of developing germ cells. Halo-tagged PGL-1 stained with an Oregon Green Halo ligand (green) and DNA with DAPI stain (blue). Images made in FIJI/ImageJ (Schindelin *et al*. 2012). **(D)** Linear diagram of *C. elegans* PGL-1. Not to scale. **(E)** Central Dogma of Molecular Biology paired with an outline of the student lab research project.

*Caenorhabditis elegans* is a simple model organism that can be modified by CRISPR-Cas9 to train inexperienced students in molecular biology and laboratory skills. The advantages of *C. elegans* include their small size for easy manipulation, transparent body for imaging, simple anatomy, ability to self-fertilize for straightforward genetics, and short life cycle. The adult hermaphrodite worm contains two large germlines with germ cells processing through cell development into oocytes **(Fig 1B)**. Sperm made in the larval stages of development is stored in the spermatheca. Oocytes cross through the spermatheca, are fertilized by sperm, and form embryos in the uterus. Despite its advantages, targeted gene editing in *C. elegans* historically has been challenging. Homologous recombination is inefficient (Plasterk and Groenen 1992; Berezikov 2004), and thus the manipulation of specific gene loci relied on forward genetic screens (Kutscher and Shaham 2014). The discovery of CRISPR-Cas9 gene editing capabilities enabled a tractable method in *C. elegans* (Dickinson *et al*. 2013; Friedland *et al*. 2013; Lo *et al*. 2013) to mutate genes and examine their phenotypes in a relatively short amount of time.

Metazoan germ cells contain discrete cytoplasmic assemblies of RNAs and proteins collectively referred to as germ granules **(Fig 1C)**. In *C. elegans*, P granules are a type of germ granule that contain specific RNAs and proteins essential for germ cell development and RNA metabolism (Phillips and Updike 2022). Proper P granule assembly is dependent on the PGL-1 scaffold protein. Structurally, PGL-1 contains an N-terminal domain (Nt), dimerization domain (DD), and a C-terminal region with RGG repeats (Ct) **(Fig 1D)** (Kawasaki *et al*. 1998; Aoki *et al*. 2016; Aoki *et al*. 2021). PGL-1 and its homologs can self-assemble into liquid condensates **(Fig 1C)** through liquid-liquid phase separation (Hyman *et al*. 2014). Little is known regarding the role of PGL-1 Ct protein region in regards to self-assembly, P granule assembly, and function in germ cell development.

This summer research module is designed to teach students the fundamentals of molecular biology through experimentation with *C. elegans* and CRISPR-Cas9 **(Fig 1E)**. The module introduces an innovating and dynamic approach that combines hands-on laboratory exposure and measurable learning assessment. In this study, students use CRISPR-Cas9 to map the protein regions in PGL-1 necessary for protein expression and P granule assembly, but the laboratory project can be adapted to any *C. elegans* and CRISPR gene editing target. Thus, this model is a practical template to teach students basic science concepts, engage students in independent laboratory research, and generate reagents for future studies.

## 2. MATERIALS AND METHODS

### 2.1 Recruitment and Assessment

Students were recruited to the lab via high school and undergraduate summer research programs at the Indiana University School of Medicine (IUSM). These programs included the Indiana University – Purdue University Indianapolis (IUPUI) Life Health Science Internship (LHSI), Indiana University Simon Comprehensive Cancer Center (IUSCCC), Indiana Medical Scientist/Engineer Training Program’s Undergraduate Summer Research Program (MSTP-USRP), and Indiana Clinical and Translational Sciences Institute (CTSI) summer research program. The applicants were chosen by their specific programs, and in most cases had the opportunity to indicate their scientific interests. Final matches were made dependent on these interests and lab availability. A Qualtrics (Qualtrics XM; www.qualtrics.com) pre- and post-test and survey were offered to lab summer students over 18 years of age. Institutional Review Board (IRB) exemption was given through Indiana University Purdue University Indianapolis (#19152). The test probed the student’s knowledge base in basic molecular biology and genetics, CRISPR, and model organisms. The survey also measured an individual’s current interest in science. The pre- and post-tests were administered on the first and last day of the student’s summer research experience. The test and survey were performed unanimously without identifiers. Due to personal issues, two students left the summer program mid-way before taking the post-test and survey due to personal issues. Pre- and post-tests were scored, and the results were graphed using GraphPad Prism software and Microsoft Excel.

### 2.2 Nematode Strains and Maintenance

Nematodes were grown on Nematode Growth Medium (NGM) plates with OP50 bacteria as food source, as described previously (Brenner 1974). All strains were propagated at 20°C. Worms were outcrossed with a wildtype N2 strain.

### 2.3 CRISPR-Cas9

Trained lab members performed all CRISPR microinjections into the gonads of young adult worms (Arribere *et al*. 2014; Kim *et al*. 2014; Paix *et al*. 2015; Ghanta *et al*. 2021). The CRISPR mix included recombinant *S. pyogenes* Cas9 (Integrated DNA Technologies, IDT), tracrRNA and *pgl-1* targeting crRNAs (IDT), and repair DNA oligo (IDT). The co-conversion approach was implemented, which involves co-injection of CRISPR-Cas9 ribonucleoproteins (RNPs) targeting the *unc-58* gene, producing uncoordinated worms that have impaired locomotion (*unc* phenotype), to select and screen worm progeny modified by the CRISPR microinjection (Arribere *et al*. 2014).

### 2.4 Polymerase Chain Reaction (PCR) and DNA sequencing

F1 *unc* L4 larvae were singled onto NGM plates with OP50 bacteria and allowed to lay eggs for approximately one day. These F1 animals were then transferred into 2x worm lysis buffer (50 mM KCl, 10 mM Tris pH 8.3, 2.5 mM MgCl2, 0.01% gelatin, 0.45% NP-40, 0.45% Tween 20 detergent, 8 units/ml Proteinase K (New England Biolabs)), lysed at 60°C for 1 hour, and PCR screened to detect the desired *pgl-1* deletions. The PCR screen used Taq polymerase (NEB), dNTPs (NEB), targeted *pgl-1* primers (IDT), and the acquired worm lysis buffer as DNA template. All primers were designed on SNAPGene software (GSL Biotech LLC; snapgene.com). F1 worms that generated the expected PCR product deletion were selected and their F2 progeny singled. These singled worms were lysed and analyzed again by PCR to identify homozygous animals. Homozygous animal samples were PCR amplified with Q5 (NEB) or KOD (Sigma) polymerase with the same primers. This PCR product was PCR purified (NEB) and Sanger sequencing performed using SupraDye v3.1 (Calibre Scientific). Unincorporated dNTPs were removed from the samples with AxyDye cleanseq magnetic beads (ThermoFisher). Samples were sequenced by ACGT (www.acgtinc.com). Sequences were analyzed by SNAPgene to confirm proper editing.

### 2.5 Immunoblot

Worms were collected in 2x sodium dodecyl-sulfate polyacrylamide (SDS) sample buffer (Bio-Rad), denatured for 10 minutes at 95°C, loaded onto 12% SDS-page gels, and transferred onto PVDF membrane (Bio Rad) using a Trans-Blot Turbo Transfer System (Bio Rad). After transfer, membrane was blocked with 5% non-fat milk in PBS-T (127 mM NaCl, 2.7 mM KCL, 10 mM Na_2_HPO_4_, 1.8 mM KH_2_PO_4_, 0.1% (w/v) Tween 20 detergent) for one hour, probed with primary V5 antibody (1:500; R&D Systems Bio-techne) overnight, washed with PBS-T, and incubated in secondary Goat anti-mouse HRP antibody (1:4000; R&D Systems Bio-techne) for at least one hour. Membrane was then washed with PBS-T and developed using SuperSignal West Pico Stable Peroxide Solution (ThermoFisher) and SuperSignal West Pico Luminol Enhancer Solution (ThermoFisher). Developed blots were imaged on a ChemiDoc MP Imaging System (Bio Rad) and analyzed on its Image Lab software (Bio Rad).

### 2.6 Confocal Microscopy

Fluorescence confocal microscopy was performed using a Zeiss AxioObserverZ1 by 3i (www.intelligent-imaging.com). Adult germline images were taken using Slidebook software (Intelligent Imaging Innovations) and a 63x objective. Worms were fixed and permeabilized in 3% paraformaldehyde (PFA) followed by DNA (1:2000 DAPI) and Halo-Oregon Green (300 nM) staining in PBS-T for one hour. Worms were wash with PBS-T after fixing and staining before being placed on slides with VECTASHIELD Mounting Medium (Vector Laboratories) for imaging. All images were analyzed using Fiji image-processing package (http://fiji.sc/Fiji) (Schindelin *et al*. 2012).

### 2.7 Statistical Analyses

GraphPad Prism software was used for graphing and statistical analyses. Pairwise comparison was determined using 2way ANOVA with multiple comparisons. Statistical significance was defined as ∗p < 0.05.

## 3. RESULTS AND DISCUSSION

Research for high school and undergraduate students promotes their retention in STEM-related fields (Nagda *et al*. 1998) and enhances students’ learning experiences (Pender *et al*. 2010). Summer break is a common time to fully immerse themselves in a research experience. Therefore, a full-time, 8-week summer teaching module was created that used CRISPR gene editing and the *C. elegans* model organism as an entrée into molecular biology. CRISPR technology is commonly used in scientific laboratories and in *C. elegans* research and is currently being developed as a cancer therapy (Baylis and Mcleod 2017). The students’ education in current gene editing methods thus had direct connections to human health, a criteria for some of these biomedical summer research programs (see Methods).

This training module included:

1. An independent research project centered around CRISPR and *C. elegans* **(Fig 2)**

**Figure 2.**
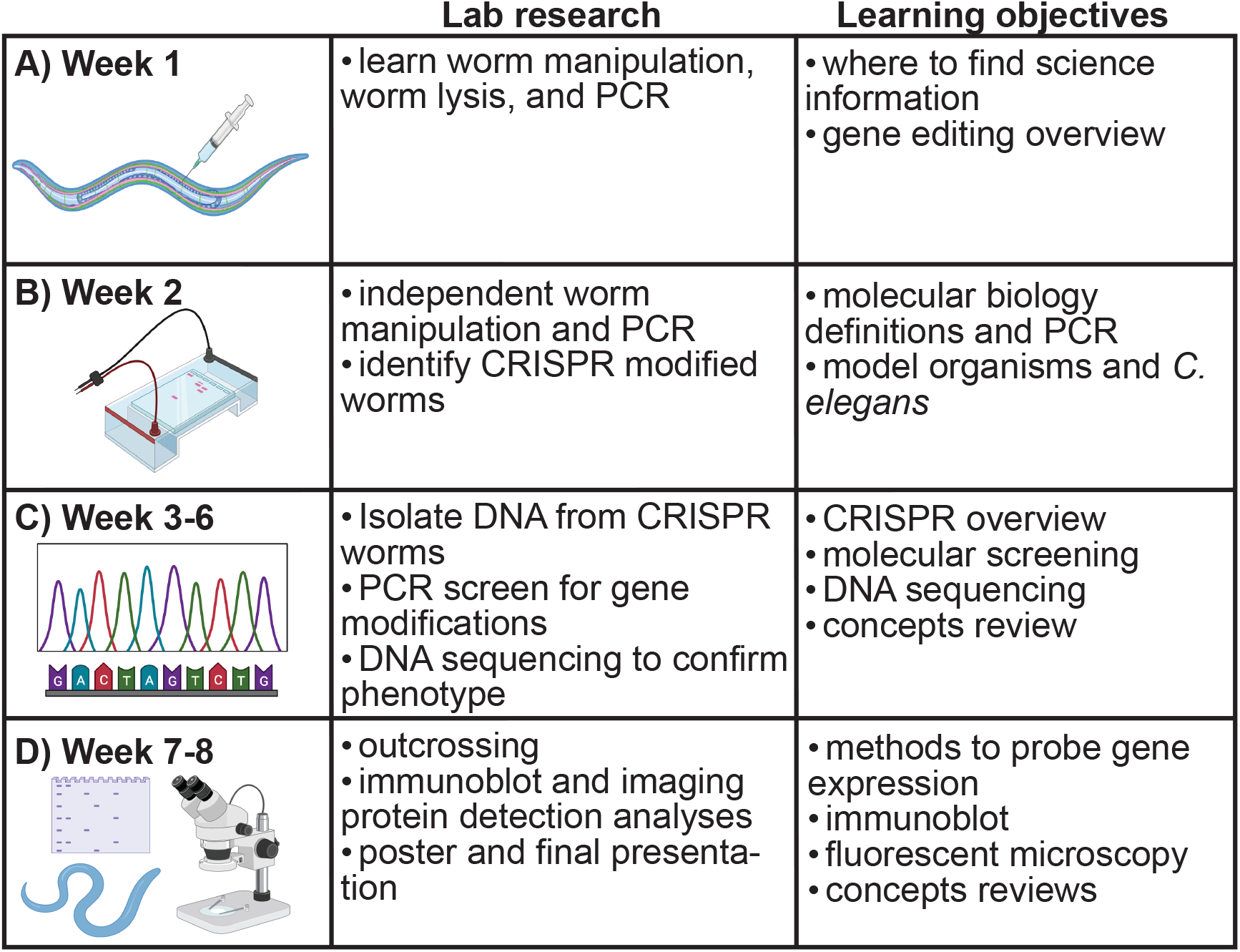
Experimental outline to identify CRISPR mutants in *C. elegans* with student learning objectives. **(A)** Week 1. Students learn to manipulate *C. elegans* and basic molecular biology methods like PCR. They also learn where to find science information and about gene editing. Adult *C. elegans* are injected with the ribonucleoprotein complex directly into their gonad. **(B)** Week 2. Students practice lab methods on their own. CRISPR modified worms are identified based on their *“unc”* phenotype and isolated onto individual plates to propagate. Students learn about molecular biology and model organisms. **(C)** Weeks 3-6. DNA is collected from *unc* worms and used to screen for genetic modifications via PCRs and gel electrophoresis. **(D)** Weeks 7-8. Once homozygous *C. elegans* with the desired mutations are identified, samples are submitted for DNA sequencing.
2. One-on-one personalized mentorship with a training mentor (e.g., graduate student)
3. Weekly wet lab assignments to provide hands-on training and step-by-step instruction toward a research project goal(s) **(Fig 2. Supplemental Table 1)**
4. Weekly dry lab assignments to provide step-by-step learning on the fundamentals of molecular biology **(Fig 2. Supplemental Table 1)**
5. Weekly hour-long molecular biology teaching and review sessions with the Lab mentor (i.e., Lab supervisor) **(Supplemental Table 1)**
6. A pre- and post-test and survey to measure students’ scientific knowledge and self-assurance **(Supplemental Table 2)**
7. End-of-the-term summer research presentation for their individual programs

The goals of the training module were to enhance students’ scientific knowledge, develop laboratory and logic skills, and explore their enthusiasm for STEM. The one-on-one mentoring enhanced communication among lab members and students and fostered meaningful interpersonal bonds between mentor and mentee. Thus, this hands-on summer research experience was tailored for the inexperienced high school and undergraduate students to teach the necessity of scientific research.

Training ran for 8 weeks, with an expected commitment of 35-40 hours per week. At the beginning of each week, students were given specific terms or questions outlined in the “dry lab” section of the summer strategic plan **(Supplemental Table 1)**. These questions focused on basic concepts in molecular biology, model organisms, CRISPR and gene editing, standard methods in DNA and protein detection, and basic laboratory techniques. Students met with their training mentors, typically a graduate student mentor, throughout the week to discuss dry lab prompts in an informal setting. This established a baseline understanding of scientific terminologies and techniques critical to the activities scheduled for the upcoming weeks. At the end of each week, a comprehensive review session was facilitated by the Lab supervisor, wherein both the dry and wet lab concepts were revisited with the students. At least one training mentor was present in the weekly reviews to ensure clarity and know what was discussed with the students. These weekly reviews enabled the mentors to gauge students’ level of comprehension and decide what to emphasize in the following weeks to fill knowledge gaps.

The student research projects used CRISPR-Cas9 gene editing to modify a gene of interest in *C. elegans*. This gene and the desired mutants were central to other research projects being concurrently pursued in the lab, with plans of using the mutant animals generated would be used in lab future research. The students in this summer cohort aimed to delete regions of *pgl-1*, a *C. elegans* gene expressed in its germline and required for proper germ cell development (Kawasaki *et al*. 1998). Prior work had determined that PGL-1 protein could conceptually be divided into Nt, DD, Ct, and RGG protein regions **(Fig 1D)** (Kawasaki *et al*. 1998; Aoki *et al*. 2016; Aoki *et al*. 2021). Students were tasked to use CRISPR-Cas-9 to delete genomic portions of *pgl-1* associated with these protein regions and test their necessity for protein expression and cell localization. All students worked with a worm strain expressing PGL-1 tagged with Halo, a modified enzyme that enabled easy labeling and protein detection (Los *et al*. 2008; Daniels *et al*. 2014; England *et al*. 2015), and a V5 epitope tag for antibody binding.

In the framework of the training module. The *C. elegans* CRISPR-Cas9 protocol was as follows:

1. CRISPR mix designed and made. CRISPR guide RNAs are designed based on the desired cleavage site and standard RNA requirements by Cas9 (Arribere *et al*. 2014; Kim *et al*. 2014; Paix *et al*. 2015; Ghanta *et al*. 2021). Recombinant Cas9 protein is incubated with commercially synthesized RNAs that target the DNA site of interest and another gene used for phenotypic screening. A repair DNA oligo is included in the mix to for the proper repair of pgl-1 and the dominant mutation of a co-injection target for a phenotype that can be used for animal screening. The *dpy-10* and *unc-58* genes were used as co-CRISPR targets, both of which are commonly used in *C. elegans* CRISPR gene editing (Arribere *et al*. 2014). All CRISPR reagents were designed and ordered by the mentors prior to the students’ arrival. The mentor also assembled the mix itself prior to use.
2. CRISPR mix injected into worms. The gonads of adult hermaphrodite, Halo-tagged PGL-1 worms were microinjected with the CRISPR mix by the mentors **(Fig 2A)**. These P_0_ parental worms were placed on single plates and incubated with food for 3-4 days until their F_1_ offspring were older larvae or adults. Before and during this incubation period, students were learning basic molecular biology and *C. elegans* methods in preparation for the subsequent steps.
3. Worms with the co-CRISPR phenotype were identified and screened. Successful gene editing of the *dpy-10* or *unc-58* co-CRISPR targets results in worms with impaired locomotion phenotype, thereby providing a distinctive phenotypic marker for the identification of CRISPR-modified worms (Arribere *et al*. 2014). Under the guidance of their mentors, students were expected to independently identify CRISPR-mutated *C. elegans* based on their unique phenotype **(Fig 2B)**, lyse worms to extract their DNA, and screen the worms by PCR analysis and gel electrophoresis **(Fig 2C)**. Modified worms were expected to have smaller amplified DNA PCR bands amplified compared to wildtype, indicating a genomic deletion at the desired site. Progeny of these worms were singled onto new plates, incubated for 1-3 days, lysed to isolate their DNA, and PCR screened again to isolate worms homozygous for the CRISPR modification.
4. Worms that were edited in the region of interest were sequenced to confirm proper repair. Once homozygous mutants were identified via PCR analysis, students independently sequenced their worms to determine whether the editing was correct. Homozygous worm DNA was PCR amplified again with a high-fidelity polymerase, and samples were sequenced by Sanger Sequencing **(Fig 2C)**. Under the mentor’s guidance, gene sequence files were aligned to the expected reference *pgl-1* genomic region to confirm proper CRISPR deletion and repair. Worms with the desired alleles were outcrossed with wildtype (N2) worms twice to lower the chances of off target CRISPR modifications. The genetic deletions were tracked by the student using PCR, as described previously.
5. Properly edited worms were analyzed by immunoblot and imaging to detect protein expression and localization **(Fig 2D)**. If the assigned *pgl-1* CRISPR deletions were successfully completed, students were given the opportunity to analyze the worm strains for protein expression and cell localization by immunoblot or imaging. N2 and Halo-tagged PGL-1 worms were used as negative and positive controls, respectively. In immunoblots, students collected adult worms in protein sample buffer, ran and SDS-PAGE gel electrophoresis, transferred the gel to a membrane, and probed the membrane for antibodies that detected the V5 epitope on Halo-tagged PGL-1 (see Methods). In imaging experiments, students collected and fixed adult worms, stained the worms with Halo ligands and DNA-binding stain, and imaged them by confocal microscopy. Thus, this experience provided students with the opportunity to learn new lab techniques, method concepts, and different perspectives on how to analyze for proteins in animals.

During the concluding week, students showcased their immersive summer experience and research endeavors through a short slideshow or poster session, required by their funding summer program **(Fig 2D)**. This allowed students to convey their scientific findings, improve their communication skills, apply critical thinking, and showcase their intellectual efforts.

The core objectives of this training module were to teach fundamental concepts in molecular biology and inspire students to think as scientists in a research laboratory setting. To evaluate the success of these goals, a pre- and post-test and survey was administered to students participating in the training module and over 18 years of age. The pre- and post-test and survey were identical to evaluate learning and growth. The test evaluated students’ knowledge of molecular biology, gene editing and CRISPR, and model organisms and *C. elegans* **(Supplemental Table 2)**. The survey measured students’ interest in science, STEM confidence, and independent learning **(Supplemental Table 2)**. The test and survey were administered at the beginning and end of the training module. Participation was optional and all results were blinded. A total of 6 students participated in the test and survey, two whom did not complete the program and thus did not take the post-evaluation.

Student testing supported the training module as a valuable strategy to teach molecular biology and instill enthusiasm for STEM research. Multiple-choice questions tested fundamental concepts in Molecular Biology, gene editing and CRISPR, and model organisms and *C. elegans* to quantitatively gauge whether students learned complicated scientific principles within a condensed period. In all three areas of study, students performed better at the end of the summer **(Fig 3A)**. The survey portion of the evaluation determined that student enthusiasm for science increased after the training module **(Fig 3B)**. Students reported an increase in their confidence to perform scientific tasks. Astoundingly, 100% of students reported substantial enjoyment for scientific learning at the end of the training module **(Fig 3B)**. Students left the training module interested in pursuing further experiences and careers in STEM. In summary, the participating students learned basic molecular biology concepts in tandem with their summer research experience and left the program interested in pursuing further STEM experiences, meeting the objectives of the program.

**Figure 3.**
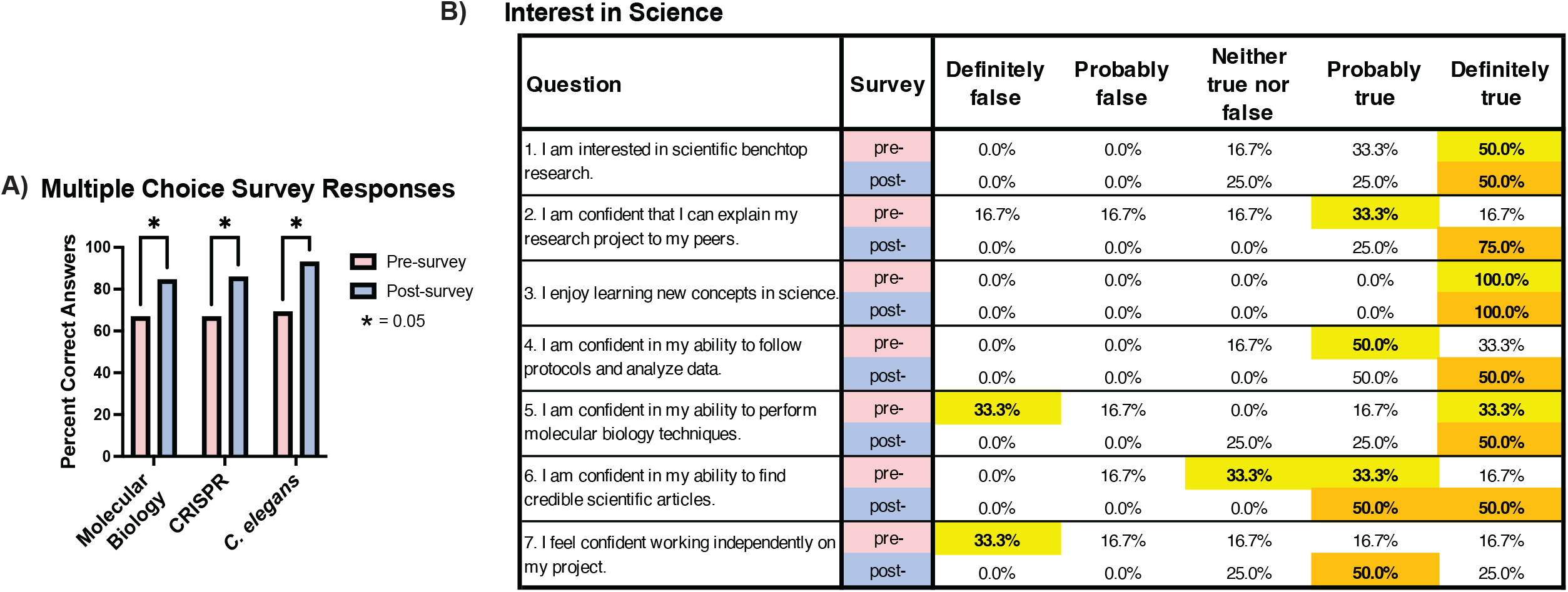
Results from pre- and post-surveys. **(A)** Table depicting the participant’s self-assurance in the training module. Between pre- and post-surveys, students reported increased confidence in all questions. Highlighted areas indicate the highest score per question. If a tie, both boxes were highlighted. **(B)** Percent correct answers for the molecular biology, CRISPR, and *C. elegans* pre- and post-survey sections. 2-way ANOVA statistical analysis was performed to compare pre- and post-survey response. Students statistically improved in all study sections.

This summer training module observed enhancements in students’ molecular biology proficiency and enthusiasm for scientific exploration. The pre- and post-surveys helped evaluate the scientific knowledge and interest gained over the experience and helped remove biases. Other studies have performed pre- and post-surveys with CRISPR study but noted variable gains in improvement. For example, an undergraduate laboratory course in CRISPR noted some RNA design concepts learned but others needing improvement (Militello and Lazatin 2017). This may be due to the differences in examination, using multiple choice in this module **(Supplemental Table 2)** versus short answer in the other study. Due to the timing and design, this study also allowed a full emersion in lab science. The students had a full work week to study the concepts and lab methods and had one-on-one mentoring. Mentors handled the advanced technical aspects of the projects, such as oligonucleotide design and CRISPR injections, while mentees were responsible for basic molecular biology tasks like PCRs, immunoblots, and DNA sequencing. This promoted a sense of teamwork and shared responsibility among participants in both the lab research and concept study. The other study was an undergraduate laboratory course and most likely could not afford the same dedication of work hours.

The opportunity to perform hands-on wet lab experiments connects molecular biology concepts with real world experience. Other studies have improved student comprehension of CRISPR-Cas9 technology solely through dry lab exposure (Pieczynski and Kee 2021). But both wet and dry lab experience has distinct benefits. Another CRISPR laboratory training study noted gains in experimental understanding but losses in data interpretation (Adame *et al*. 2016). While this may seem like a negative, it may also reflect students realizing that science requires more knowledge and training than what can just be achieved in the classroom. This work noted that students were still enthusiastic about STEM careers. Thus, combining research with learning concepts can maximize a student’s overall experience.

Overall, this training module incorporates diligent assessment methods, hands-on experience, and a collaborative learning environment that enhances science education. While the study focused on molecular biology and CRISPR gene editing, the approach can be implemented in any science topic being investigated by research laboratories accepting summer high school and undergraduate students.

## ACKNOWLEDGMENTS

The authors thank other members of the Aoki Lab for their supportive secondary mentorship with summer students and project discussions. They thank all the summer student participants. This work was supported by Indiana University – Purdue University Indianapolis (IUPUI) Life Health Science Internship (LHSI), Indiana University Simon Comprehensive Cancer Center (IUSCCC), Indiana Medical Scientist/Engineer Training Program’s Undergraduate Summer Research Program (MSTP-USRP), and Indiana Clinical and Translational Sciences Institute (CTSI) summer research program. S.T.A. is funded by the NIH/NIGMS (R35 GM142691) and received start-up funds from the Indiana University School of Medicine and its Precision Health Initiative (PHI).

## 6. SUPPLEMENTARY INFORMATION

### 6.1 Supplemental Table 1. Summer strategic plan

**Supp. Table 1.** Summer stragetic plan detailing dry and wet lab assignments.

| Week | Dry Lab | Wet Lab |
| --- | --- | --- |
| 1 | a) How do you find science information? Name several (3) strategies for finding information from home.<br>b) False information in science. How can you tell real, science-based information from fake information?<br>c) What is gene editing? Name and describe types of gene editing.<br>d) What can gene editing be used for? Describe an example used in medicine.<br>e) Friday Recap | a) Lab tour and safety<br>b) Discuss lab duties (dishware)<br>c) Practice worm picking<br>d) Perform worm lysis<br>e) PCR set up/protocol<br>f) Perform PCR gel electrophoresis |
| 2 | a) Define DNA, RNA, proteins, chromosomes, genes, alleles, cells, organelles, nucleus, animals, genotype, phenotype, strain, allele.<br>b) What are model organisms? Name common ones used in research. What are they used for?<br>c) What is *Caenorhabditis elegans*?<br>d) When was it discovered?<br>e) What information has *C. elegans* given us about biology since it's discovery?<br>f) What is PCR? How does it work? How can we use it to identify animal strains?<br>g) Friday Recap | a) Continue worm lysis<br>b) Continue PCR<br>c) Continue PCR gel electrophoresis<br>d) Perform primer design & screening<br>e) [CRISPR injections] |
| 3 | a) What is CRISPR? What can we use it for? How is it better than other gene editing methods?<br>b) How does CRISPR work? Describe the general mechanism. What components are necessary for the reaction?<br>c) How is CRISPR done in *C. elegans*?<br>d) When designing a CRISPR guide targeting a specific gene, what criteria are necessary for the target?<br>e) Designing the desired allele. What can we insert or change in the animal's genome to study basic questions in biology?<br>f) Design a screening method for CRISPR. How can we identify our repair of interest?<br>g) Friday Recap | a) Worm phenotype screening (uncs)<br>b) Continue worm lysis<br>c) Continue PCR<br>d) Continue PCR gel electrophoresis |
| 4/5 | a) How do you screen worms for modifications?<br>b) What is Sanger sequencing? How does it work?<br>c) DNA sequencing and sequencing analyses<br>d) Friday Recap | a) Worm phenotype screening (uncs)<br>b) Continue worm lysis<br>c) Continue PCR<br>d) Continue PCR gel electrophoresis<br>a) Worm phenotype screening (homozygous)<br>b) Continue worm lysis<br>c) Continue PCR<br>d) Continue PCR gel electrophoresis<br>e) DNA Sanger sequencing |
| 6/7 | a) Testing our CRISPR'd animal: molecular biology and biochemical assays (Imaging/Immunoblot)<br>b) How is protein different from DNA or RNA? What assays can we use to detect our modified protein?<br>c) Confocal Imaging, Immunoblot, and CRISPR review.<br>d) Poster outline and drafting<br>e) Friday Recap | a) Worm outcrossing<br>b) Review immunoblot protocols<br>c) Review confocal imaging protocols |
| 8 | a) Summer wrap-up<br>b) Practice presentation<br>c) Formal presentation<br>d) Friday Recap and Final Evaluation | a) Finish experiments<br>b) Freeze worms<br>c) Lab clean up |

### 6.2. Supplemental Table 2. Pre- and post-survey

**Supp. Table 2.** pre- and post- survey sections with corresponding questions and answers.

| Questionnaire |  |  |  |
| --- | --- | --- | --- |
| SECTION | QUESTION | OPTIONS | ANSWER |
| A) INTEREST IN SCIENCE | Rate each statement on a scale of 1 to 5 (1 being not confident/not interested and 5 being very confident/very interested) | 1. I'm interested in scientific benchtop research.<br>2. I'm confident that I can explain my research project to my peers.<br>3. I enjoy learning new concepts in science.<br>4. I'm confident in my ability to follow protocols and analyze data.<br>5. I'm confident in my ability to perform molecular biology techniques.<br>6. I'm confident in my ability to find credible scientific articles.<br>7. I feel confident working independently on my project. | N/A |
| B) MOLECULAR BIOLOGY | 1. What is the central dogma of molecular biology? | a) Process that explains the flow of genetic information, from DNA to functional protein<br>b) Theory explaining cellular replication in mammals<br>c) Physical process that explains protein folding | A |
|  | 2. What is RNA? | a) Nucleic acid present in all living cells responsible for the regulation and expression of genes<br>b) Large biomolecules comprising of one or more long chains of amino acid residues<br>c) Deoxyribonucleic acid is a polymer composed of two polynucleotide chains that coil around each other to form a double helix carrying genetic instructions for the development, functioning, growth | A |
|  | 3. What is RNA used for in cells? | a) To make proteins<br>b) As an enzyme<br>c) To regulate other RNAs<br>d) All of the above | D |
|  | 4. What is a gene? | a) Proteins that act as biological catalysts to accelerate chemical reactions<br>b) Minute particles consisting of RNA and associated proteins that perform biological protein synthesis<br>c) Basic unit of heredity and a sequence of nucleotides in DNA | C |
|  | 5. What is protein translation in biology? | a) Process by which a cell makes proteins using the genetic information carried in mRNA<br>b) Process in which a gene's DNA sequence is copied to make an RNA molecule<br>c) Process by which RNA strands become longer due to the addition of new nucleotides | A |
|  | 6. What are the steps in PCR? | a) Denaturation, annealing, and extension<br>b) Protein transfer, block, and probe<br>c) Paraformaldehyde fix, PBS wash, and stain | A |
|  | 7. What are the components of PCR? | a) Filter paper, transfer buffer, and a nitrocellulose or PVDF membrane<br>b) DNA, primers, free nucleotides, and polymerase<br>c) SDS, b-mercaptoethanol (BME), bromophenol blue, glycerol, and Tris-glycine | B |
|  | 8. DNA gel electrophoresis separates DNA fragments dependent on their...? | a) Sequence composition<br>b) Size<br>c) Folding | B |

| Questionnaire Cont. |  |  |  |
| --- | --- | --- | --- |
| SECTION | QUESTION | OPTIONS | ANSWER |
| C) CRISPR | 1. What is CRISPR? | a) Technology for freezing worms<br>b) Technology for gene editing<br>c) Technology for protein extraction | B |
|  | 2. What does CRISPR stand for? | a) Charpentier-Ressa-Inman Systemic Process for Recombination<br>b) Computed Ribosomal Interspaced Soluble Patches for Recombination<br>c) Clustered Regularly Interspaced Short Palindromic Repeats | C |
|  | 3. Which of the following use the CRISPR/Cas system naturally? | a) Eukaryotes<br>b) Prokaryotes<br>c) Viruses | B |
|  | 4. What is the enzymatic purpose of the Cas9 protein? | a) To elongate DNA<br>b) To connect two DNA fragments<br>c) To cut DNA | C |
|  | 5. What are the components in CRISPR-Cas9? | a) Guide RNA and enzyme<br>b) Guide DNA and enzyme<br>c) Repair DNA and enzyme | A |
|  | 6. What is a ribonucleoprotein complex? | a) Complexes of DNA and protein<br>b) Multiple proteins assembled in a complex<br>c) Complexes of RNA and protein | C |
|  | 7. Which technique to screen the genome for CRISPR modifications in *C. elegans*? | a) Imaging<br>b) PCR<br>c) Immunoblot | B |
| D) *C. elegans* | 1. What is a model organism? | a) Non-human species that are used in the laboratory to help scientists understand biological processes<br>b) Human tissue samples that are used in the laboratory to help scientists understand biological processes<br>c) Cells grown under controlled conditions, generally outside their natural environment that are used in the laboratory to help scientists understand biological processes | A |
|  | 2. What are examples of model organisms? | a) Human<br>b) Bone fragment<br>c) Nematode worm | C |
|  | 3. What characteristics of *C. elegans* make it a great model organism in research laboratories? | a) Small in size<br>b) Quick reproductive life cycle<br>c) Translucent<br>d) All of the above<br>e) None of the above | D |
|  | 4. What is a major advantage of *C. elegans* versus other model organisms? | a) *C. elegans* has an advanced endocrine system<br>b) *C. elegans* can undergo binary fission<br>c) *C. elegans* can self-fertilize | C |
|  | 5. *C. elegans* can naturally be found in: | a) Water<br>b) Laboratories<br>c) Soil | C |
|  | 6. Germ cells make: | a) Connective tissue<br>b) Internal organs<br>c) Gametes | C |
|  | 7. To obtain modified CRISPR-Cas9 worms, we modify an organism's: | a) Germline<br>b) Intestine<br>c) Neurons | A |

## Notes

### Competing Interest Statement

The authors have declared no competing interest.

